# HMGB1 acts as an agent of host defense at the gut mucosal barrier

**DOI:** 10.1101/2023.05.30.542477

**Authors:** A-M C Overstreet, B Anderson, M Burge, X Zhu, Y Tao, CM Cham, B Michaud, S Horam, N Sangwan, M Dwidar, X Liu, A Santos, C Finney, Z Dai, VA Leone, JS Messer

**Affiliations:** Lerner Research Institute, Cleveland Clinic, Cleveland OH, United States; Department of Medicine, University of Chicago, Chicago IL, United States; Department of Gastroenterology, Shanghai Renji Hospital, Shanghai, China; Department of Animal & Dairy Sciences, University of Wisconsin-Madison, Madison WI, United States

## Abstract

Mucosal barriers provide the first line of defense between internal body surfaces and microbial threats from the outside world.^1^ In the colon, the barrier consists of two layers of mucus and a single layer of tightly interconnected epithelial cells supported by connective tissue and immune cells.^2^ Microbes colonize the loose, outer layer of colonic mucus, but are essentially excluded from the tight, epithelial-associated layer by host defenses.^3^ The amount and composition of the mucus is calibrated based on microbial signals and loss of even a single component of this mixture can destabilize microbial biogeography and increase the risk of disease.^4–7^ However, the specific components of mucus, their molecular microbial targets, and how they work to contain the gut microbiota are still largely unknown. Here we show that high mobility group box 1 (HMGB1), the prototypical damage-associated molecular pattern molecule (DAMP), acts as an agent of host mucosal defense in the colon. HMGB1 in colonic mucus targets an evolutionarily conserved amino acid sequence found in bacterial adhesins, including the well-characterized Enterobacteriaceae adhesin FimH. HMGB1 aggregates bacteria and blocks adhesin-carbohydrate interactions, inhibiting invasion through colonic mucus and adhesion to host cells. Exposure to HMGB1 also suppresses bacterial expression of FimH. In ulcerative colitis, HMGB1 mucosal defense is compromised, leading to tissue-adherent bacteria expressing FimH. Our results demonstrate a new, physiologic role for extracellular HMGB1 that refines its functions as a DAMP to include direct, virulence limiting effects on bacteria. The amino acid sequence targeted by HMGB1 appears to be broadly utilized by bacterial adhesins, critical for virulence, and differentially expressed by bacteria in commensal versus pathogenic states. These characteristics suggest that this amino acid sequence is a novel microbial virulence determinant and could be used to develop new approaches to diagnosis and treatment of bacterial disease that precisely identify and target virulent microbes.

## Main

The acellular mucosal barrier lies over the colon and normally limits physical and biochemical contact between microbes and the host epithelium.^8^ When this first barrier fails, microbes adhere to intestinal epithelial cells (IEC) leading to high levels of toxins and pro-inflammatory microbial components at the IEC surface, invasion into IEC, and translocation across the cellular barrier into deeper tissues.^9–11^ Intestinal epithelial cells are responsible for production and transport of acellular barrier components into the gut lumen and form the cellular portion of the barrier.^12^ We previously demonstrated that mice conditionally deficient in IEC HMGB1 (ΔIEC) are highly susceptible to chemical and immune-mediated colitis.^13^ However, we also observed high levels of HMGB1 in IEC of healthy humans and mice along with evidence of cytokine abnormalities in mice lacking IEC HMGB1, even when unchallenged.^13^ These factors led us to investigate physiologic functions of HMGB1 at the colonic mucosal barrier.

## HMGB1 is released into colonic mucus in response to the gut microbiota

HMGB1 was highly concentrated at the luminal surface of the mouse colon in the tight, epithelial-associated mucus layer (**Fig. 1a, Extended data Fig. 1**). In littermate ΔIEC mice, HMGB1 was absent from IEC bodies and profoundly decreased in mucus, suggesting that IEC are the primary source of HMGB1 in colonic mucus (**Fig. 1b, c**). HMGB1 protein was likewise detectable in IEC from C57BL/6 germ-free (GF) mice and the levels appeared only mildly decreased compared to specific pathogen free (SPF) C57BL/6 mice (**Fig. 1d, e**). However, the presence of HMGB1 in the gut lumen was dependent on microbiota as HMGB1 staining was absent from the colon surface and HMGB1 protein was undetectable in stool from GF mice (**Fig. 1d, f**). HMGB1 was assayed in stool because GF mice produce very little colonic mucus.^5, 14^ Taken together, these data show that HMGB1 is released from IEC into the colonic mucus in response to the gut microbiota.

**Figure 1.**
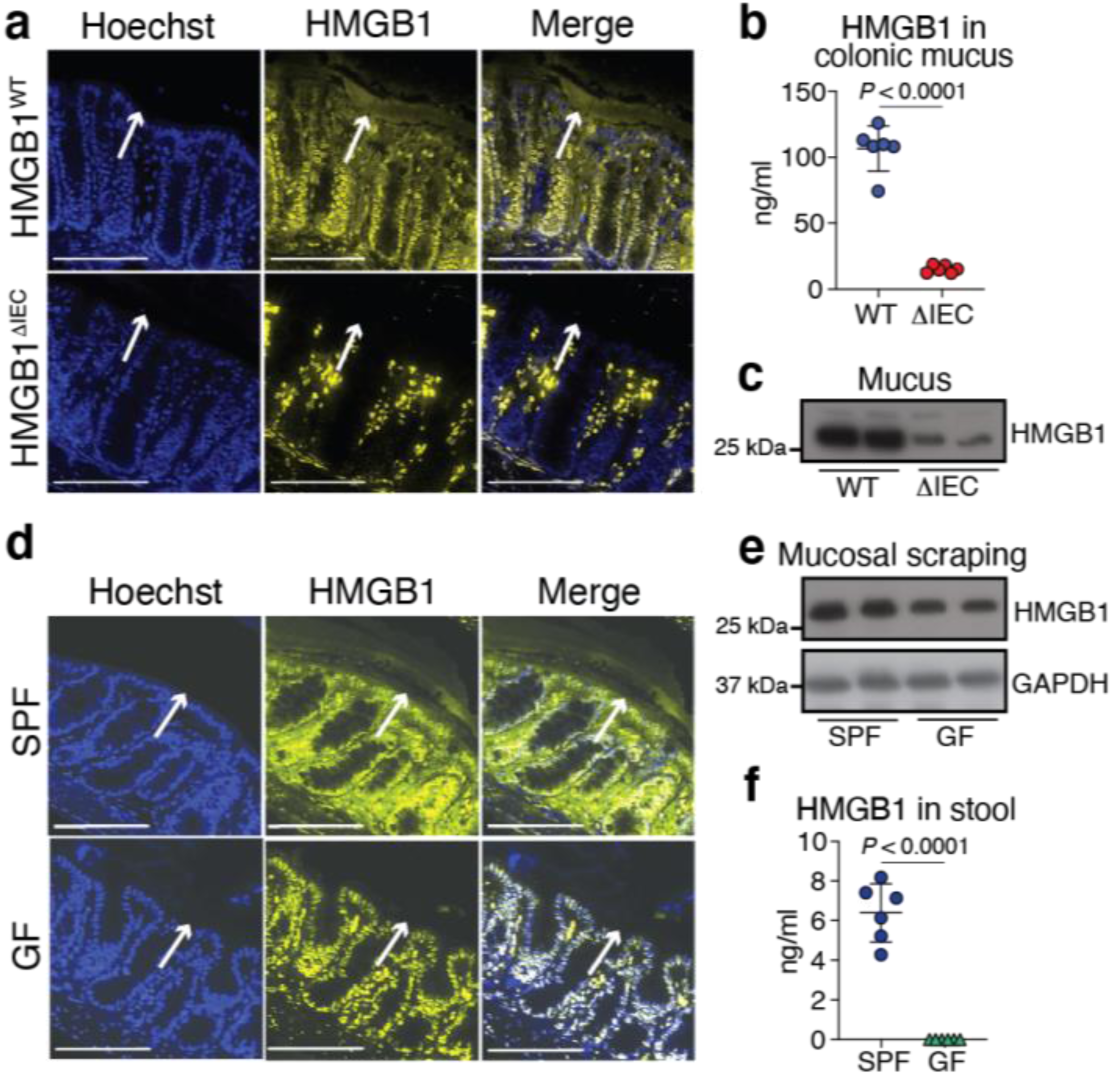
HMGB1 is released into colonic mucus in response to the gut microbiota. **a,** Immunostaining for HMGB1 (yellow) and bisbenzimide H 33258 (Hoescht, blue) in Carnoy’s fixed proximal colon sections from HMGB1^WT^ and HMGB1^ΔIEC^ mice. Arrows indicate the epithelial surface. (n=20) **b,** HMGB1 concentration in colonic mucus from HMGB1^WT^ and HMGB1^ΔIEC^ mice measured by ELISA. (n=6) **c,** Immunoblotting for HMGB1 in colonic mucus from HMGB1^WT^ and HMGB1^ΔIEC^ mice. (n=7) **d,** Immunostaining for HMGB1 (yellow) and Hoescht (blue) in Carnoy’s fixed proximal colon sections from SPF and GF C57BL/6 mice. Arrows indicate the epithelial surface. (n=6) **e,** Immunoblotting for HMGB1 in mucosal scrapings from SPF and GF C57BL/6 mice. (n=4) **f,** HMGB1 concentration in stool from SPF and GF C57BL/6 mice measured by ELISA. (n=6) Data are mean ± s.d. Significance determined by Student’s two-tailed T-tests. Each datapoint represents one individual mouse. Scale bars, 100 µm. Original magnification 400x

## HMGB1 prevents bacterial invasion into the inner mucus layer of the colon

The presence of HMGB1 in the gut lumen suggested that it could affect the gut microbiota, so we examined the influence of HMGB1 on the composition and behavior of the gut bacterial community. The normal colon boasts a diverse, abundant microbial community that is physically separated from the epithelial surface by the inner mucus barrier.^8^ In mice lacking mucosal HMGB1, this physical separation was essentially lost leading to close proximity of the microbial community to host tissue along with an increase in bacterial DNA associated with host tissue (**Fig. 2a, b, c**). The change in microbial biogeography was not due to a decrease in mucus production in ΔIEC mice (**Extended data Fig. 2a, b, c**). HMGB1 labeled microbes in the gut lumen, leading us to assess whether HMGB1 has direct effects on the microbiota (**Extended data Fig 3a**). The mucosal-associated microbial community did differ between WT and ΔIEC mice (**Extended data Fig. 3b, c**). Most notably, the taxonomic distinction normally present between stool and mucosal-associated bacteria was diminished in ΔIEC mice, reinforcing that HMGB1 is responsible for excluding microbiota from the inner mucus layer (**Extended data Fig. 3d**). However, taxonomic differences in mucosal-associated bacteria between the mouse genotypes were primarily appreciated at the strain level, suggesting that HMGB1 does not exert strong selection pressure on gut bacteria (**Extended data Fig. 3e**).

**Figure 2.**
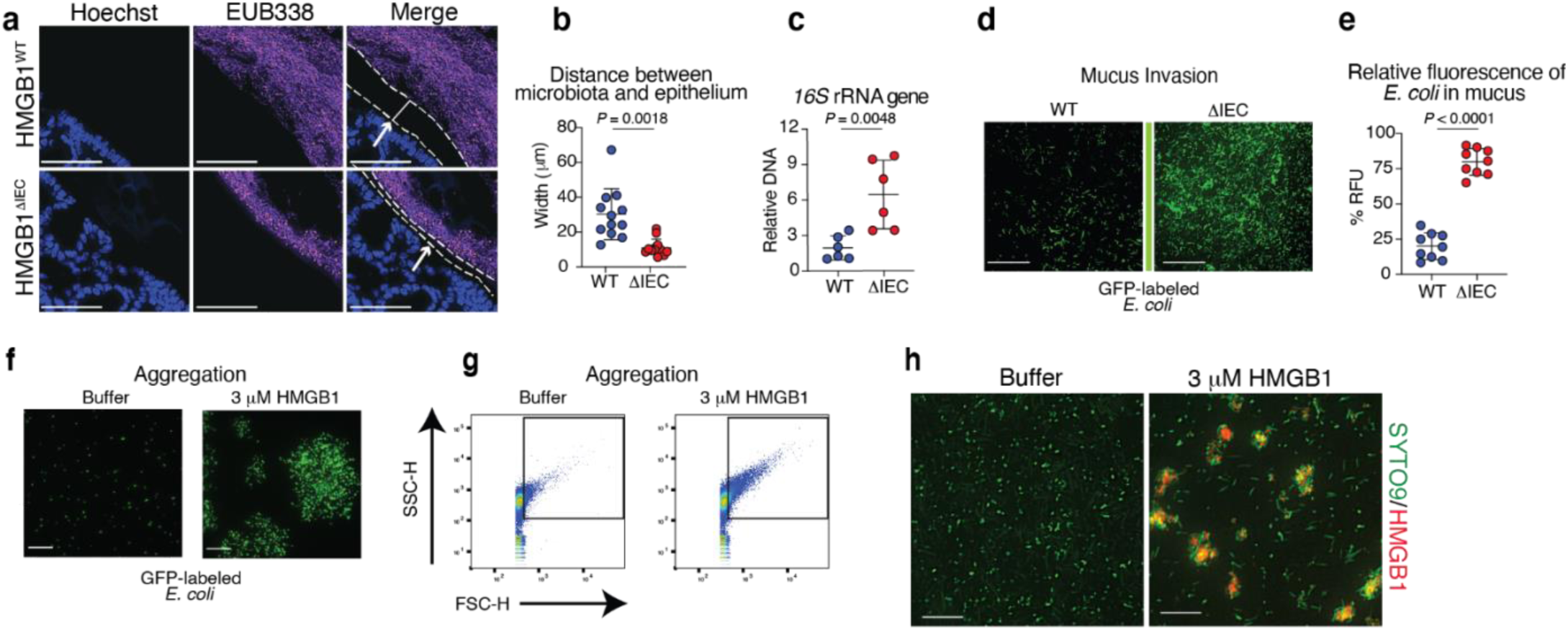
HMGB1 prevents bacterial invasion into the inner mucus layer of the colon. **a,** Fluorescence *in situ* hybridization (FISH) using the EUB338 probe (purple) and Hoescht (blue) in Carnoy’s fixed proximal colon sections from HMGB1^WT^ and HMGB1^ΔIEC^ mice. Arrows indicate the epithelial surface. Dotted lines are placed at the epithelial surface and inner edge of the microbial community. Straight line indicates distance between host tissues and microbial community. (n=12) **b,** Distance measured between epithelium and bacterial cells in images represented in (a). Each datapoint is an average of 5 measurements for one individual mouse. **c,** Quantitative PCR for the bacterial 16S rRNA gene in 1 cm of colonic tissue from HMGB1^WT^ and HMGB1^ΔIEC^ mice. (n=6) **d,** Invasion of green fluorescent protein (GFP) labeled *E. coli* (SWW33) into mucus isolated from HMGB1^WT^ or HMGB1^ΔIEC^ mice. Original magnification 200x (n=3; 3 replicates) **e,** Percentage of the total GFP signal in mucus from HMGB1^WT^ vs. HMGB1^ΔIEC^ mice in images represented in (d). (n=3; 3 replicates) **f,** Appearance of GFP labeled *E. coli* (SWW33) exposed to buffer (control) or HMGB1. (n=3; 3 replicates) **g,** Flow cytometry of aggregates in samples of GFP labeled *E. coli* (SWW33) exposed to buffer (control) or HMGB1. (n=3; 3 replicates) **h,** Appearance of SYTO 9 labeled microbiota from C57BL/6 mice exposed to buffer (control) or HMGB1 labeled with AF647. Scale bars, 20 µm (n=3; 3 replicates) Data are mean ± s.d. Significance determined by Student’s two-tailed T-tests. Each datapoint represents one individual mouse. Scale bars, 100 µm and original magnification 400x, unless otherwise noted.

We considered that loss of HMGB1 might allow normally commensal microbes to penetrate colonic mucus. Mucus is thought to primarily act as a physical anti-microbial barrier in the colon.^8^ The most abundant protein in intestinal mucus, Mucin 2 (Muc2), is a large, heavily glycosylated protein that oligomerizes to form a dense meshwork that blocks bacterial movement.^15^ Additionally, oligosaccharides attached to Muc2 are the same oligosaccharides that bacterial adhesins bind to on the surface of host cells, so they serve as decoy adhesion sites and arrest movement through mucus.^16, 17^ Microbial invasion into mucus was limited when HMGB1 was present in mucus (**Fig. 2d, e**). *E. coli* was chosen for these studies because it is a well-characterized gut commensal organism that has been associated with colitis in animal models and in human inflammatory bowel disease (IBD) patients.^18^ Exposure to HMGB1 also caused *E. coli* and complex microbiota to aggregate, similar to what has been reported for secretory immunoglobulin A (sIgA) in small intestinal mucus (**Fig. 2f, g, h**).^19^ Aggregation by sIgA traps bacteria that enter small intestinal mucus and blocks their access to adhesion targets on host epithelial cells.^20^ Our findings suggest that HMGB1 is performing a similar function in colonic mucus. Microbes coming into contact with HMGB1 are aggregated and prevented from migrating through the mucus and interacting with the epithelial surface of the colon.

## HMGB1 binds and inactivates the bacterial adhesin FimH through an evolutionarily conserved amino acid sequence

The observation that HMGB1 binds directly to gut microbes *in vivo* and *in vitro* led us to hypothesize that HMGB1 targets one or more proteins expressed on the surface of bacteria. In our previous work, we identified an amino acid sequence shared between the mammalian proteins Beclin-1 and Autophagy protein 5 (Atg5) that was targeted by cytosolic HMGB1 during microbial stress.^13^ Querying the PROSITE database using a motif derived from this sequence yielded a large number of hits in known or predicted bacterial adhesins, including FimH, a component of type 1 fimbrial (T1F) adhesion and perhaps the best characterized bacterial adhesin (**Fig. 3a**).^21, 22^ FimH is a phase-inducible virulence factor that is carried by Enterobacteriaceae, including *E. coli*, and has been implicated in infectious diarrheal diseases, urinary tract infections, extraintestinal infections, colorectal cancers, and inflammatory bowel diseases.^22–27^ Expression of FimH is low in *E. coli* when they are in a commensal state, but it is highly expressed by pathogenic *E. coli* strains or when commensal strains become virulent.^28, 29^ We first verified HMGB1 interaction with the conserved amino acid motif target <u>o</u>f <u>H</u>MGB<u>1</u> (ToH1) in FimH expressed by *E. coli*. Recombinant human HMGB1 (rHMGB1) bound to *E. coli* expressing WT FimH, but the numbers of bacteria positive for HMGB1 and the amount of HMGB1 bound to each individual bacterium were significantly lower when cells lacked FimH (ΔFimH) or expressed FimH mutated in ToH1 (ΔFimH^MUT)^ (**Fig. 3b, c and Extended data Fig. 4a, b, c, d**). We next employed a label transfer assay to interrogate direct protein-protein interaction between HMGB1 and the IEC binding lectin domain of FimH (FimH_LD_). Label transferred to WT rFimH_LD_, but not rFimH_LD_ mutated in ToH1, demonstrating that HMGB1 binds to FimH through ToH1 (**Fig. 3d**). Inclusion of mannose, the ligand bound by FimH for T1F adhesion, did not alter the efficiency of label transfer from rHMGB1 to WT rFimH. This and published structural data showing that ToH1 is distinct from the mannose binding site in FimH indicate that HMGB1 does not compete with mannose for binding to FimH.^30^ Auto transfer of label from rHMGB1 to itself was also observed in the WT rFimH_LD_ reactions, supporting that HMGB1 forms an oligomer, most likely a dimer, when it interacts with FimH. This is consistent with the ability of HMGB1 to bind to FimH on more than one bacterial cell and cause aggregation, with literature reporting that HMGB1 dimerizes, and with concentrations of the two proteins that allow for maximal label transfer.^31^ We then examined whether HMGB1 could limit *E. coli* adhesion to host cells. rHMGB1 inhibited *E. coli* adhesion to IEC at a level that was equivalent to mannose and inhibited red blood cell (RBC) agglutination in a dose-dependent manner (**Fig. 3e and Extended data Fig. 5**). rHMGB1 also inhibited binding between rFimH_LD_ and mannose showing that reduction in adhesion occurs due to the direct interaction between HMGB1 and FimH (**Fig. 3f**). Mutation of ToH1 in FimH severely impaired RBC agglutination by *E. coli*, implying that this sequence is necessary for the adhesin function of this protein (**Fig. 3g**).

**Figure 3.**
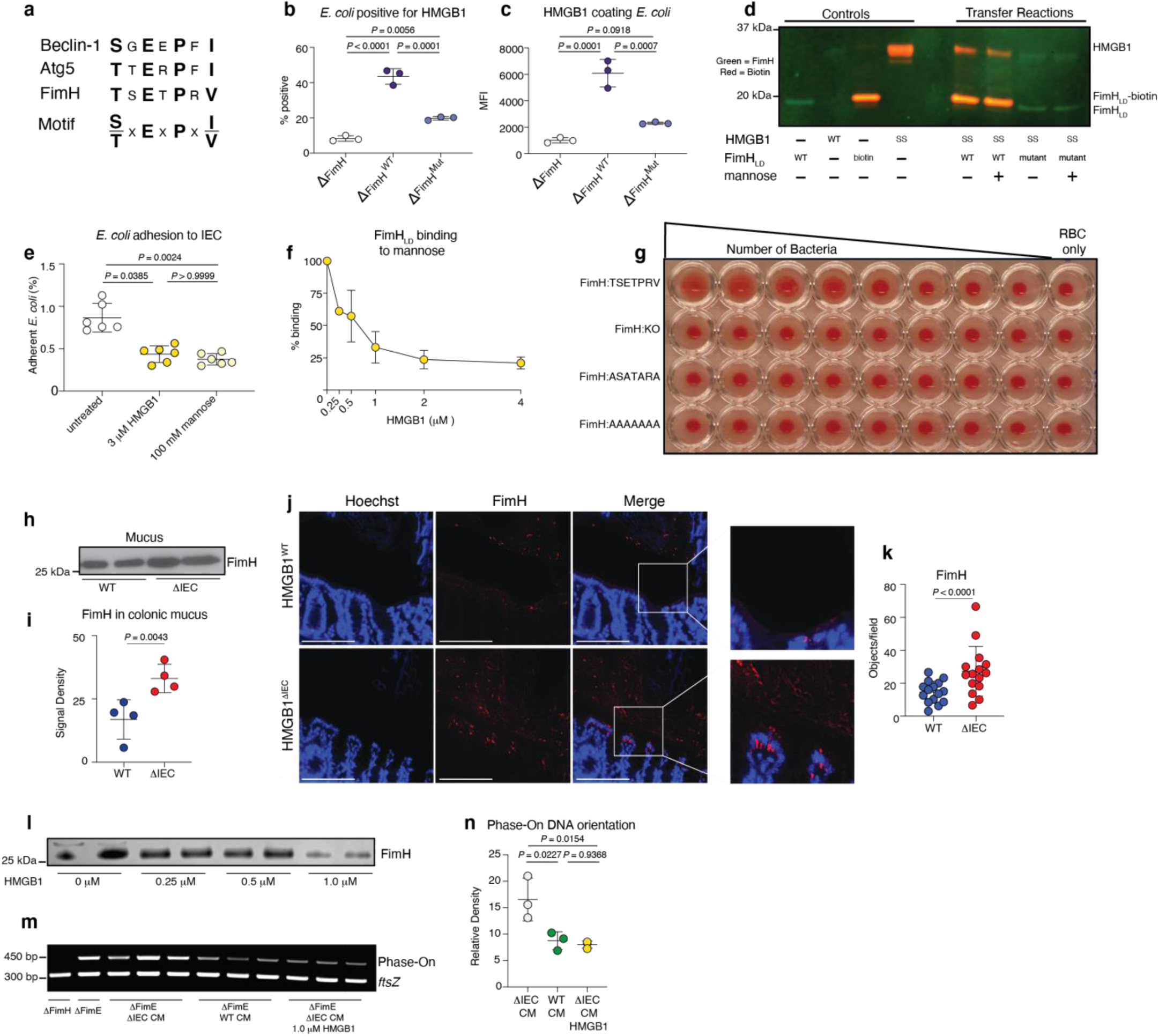
HMGB1 binds, inactivates, and regulates expression of the bacterial adhesin FimH through an evolutionarily conserved amino acid sequence. **a,** Amino acid sequence similarities among the known human HMGB1 target proteins Beclin-1 and Atg5 and bacterial FimH. The putative HMGB1 interaction motif was derived from the amino acid sequence similarities between Beclin-1 and Atg5 and common amino acid replacements. **b,c,** Flow cytometry of rHMGB1 binding to *E. coli* (BW25113) knocked out for FimH (ΔFimH) or ΔFimH *E. coli* complemented with plasmids encoding wild type FimH (ΔFimH^WT^; TSETPRV) or FimH mutated in the conserved residues of the putative motif (ΔFimH^MUT^; ASATARA). Both the percent *E. coli* positive for HMGB1 (b) and the amount of HMGB1 protein (mean fluorescence intensity (MFI)) bound to each bacterium (c) were assessed. (n=3; 3 replicates) **d,** Label transfer of Sulfo-SBED from HMGB1 to recombinant FimH lectin domain (FimH_LD_). Recipient proteins were wild type FimH_LD_ (WT; TSETPRV) or FimH_LD_ mutated in the conserved amino acid residues (mutant; ASATARA) of the putative interaction motif. Transfer was assessed in the absence or presence of mannose. (3 replicates) e, *E. coli* colony forming units (CFU) adherent to Caco2 IEC as a percent of input. *E. coli* were treated with buffer, rHMGB1, or mannose prior to addition to the IEC. (n=6; 3 replicates) **f,** Percentage of FimH_LD_ protein bound to a mannose coated plate in the presence of increasing amounts of rHMGB1. (n=3; 3 replicates) **g,** RBC agglutination by *E. coli* (SWW33) expressing wild type FimH (FimH:TSETPRV), knocked out for FimH (FimH: KO), or expressing FimH mutated in the ToH1 sequence (FimH:ASATARA and FimH:AAAAAAA). Numbers of bacteria decrease from left to right. (3 replicates) **h,** Immunoblotting for FimH in mucus isolated from HMGB1^WT^ and HMGB1^ΔIEC^ mice. (n=4) **i,** Quantification of band densitometry of immunoblots represented in (h). (n=4) **j,** Immunostaining for FimH (red) and Hoescht (blue) in Carnoy’s fixed proximal colon sections from HMGB1^WT^ and HMGB1^ΔIEC^ mice. Scale bars, 100 µm. Original magnification 400x. (n=15) **k,** Quantification of FimH positive bacteria in images represented in (j). **l,** Immunoblotting for FimH in *E. coli* (SWW33) exposed to increasing amounts of HMGB1. (3 replicates) **m,** PCR determination of the orientation of the DNA switch region govering Fim gene expression in *E. coli* (ΔFimE) treated with media conditioned by IEC organoids derived from HMGB1^ΔIEC^ mice (ΔIEC CM), HMGB1^WT^ mice (WT CM), or ΔIEC CM supplemented with rHMGB1. Phase-on denotes switch oriented toward production of Fim genes. ftsZ is used for normalization. **n,** Relative band density of phase-on in (l). Data are mean ± s.d. Significance determined by Student’s two-tailed T-tests for pairwise comparisons or one-way ANOVA with a Tukey post hoc test. Each datapoint represents one individual mouse.

## HMGB1 regulates expression of type 1 fimbrial adherence machinery in *E. coli*

In mice lacking IEC HMGB1, FimH protein levels were higher in colonic mucus and in tissue than WT littermates (**Fig. 3h, i, j, k and Extended data Fig. 6a, b, c**). This is consistent with our data demonstrating overall increased numbers of bacteria at the epithelial surface in ΔIEC mice. However, the increase in FimH could be due to increased numbers of bacteria expressing the same amount of FimH, increased FimH expression on each bacterium without a change in the number of bacteria, or both. We did not appreciate differences in relative abundance of *E. coli* or Enterobacteriaceae between the mouse genotypes by 16s rRNA gene sequencing and *in vitro* assays did not identify an effect of HMGB1 on *E. coli* colony forming units (**Extended data Fig. 6d**). Therefore, we considered that HMGB1 could regulate bacterial expression of FimH. Exposure to rHMGB1 decreased FimH expression by commensal *E. coli* in a dose-dependent manner. (**Fig. 3l**). In order to determine whether HMGB1 causes *E. coli* to turn off FimH expression or prevents bacteria from turning it on, we took advantage of a FimE knockout (ΔFimE) strain of *E. coli*.^32^ The genes required for type 1 fimbriae are arranged in a single operon that is regulated by a DNA switch region.^33^ The switch between fimbrial producing (FimON) and non-producing (FimOFF) states is regulated by two recombinases, FimB and FimE.^33^ When FimE is knocked out, bacteria can switch to FimON, but switching to FimOFF is impaired, providing a relative “counter” for the switch to FimON.^33^ Media conditioned by WT primary intestinal epithelial cells repressed FimON in comparison to media conditioned by cells derived from ΔIEC mice (Fig. 3m, n). Furthermore, addition of HMGB1 to the ΔIEC conditioned media repressed FimON similarly to WT media. Thus, these combined results indicate that HMGB1 in the environment binds to its evolutionarily conserved target sequence (ToH1) in FimH, inhibits FimH binding to its target ligand mannose, impairs T1F adhesion to host cells, and suppresses expression of this adhesion mechanism.

## HMGB1 mucosal defense is compromised in human ulcerative colitis

We previously reported intracellular HMGB1 deficiency in IEC from patients with IBD.^13^ Our discovery of HMGB1 mucosal defense released from IEC led us to suspect that this defense could also be compromised in IBD patients. There are two major subtypes of IBD, Crohn’s disease and ulcerative colitis.^34^ Ulcerative colitis affects only the colon and is thought to result from tissue damage initiated at the luminal surface of the epithelium.^35^ Surface-associated HMGB1 was generally easily appreciable in resected colon tissue from non-IBD patients, whereas tissue from UC patients commonly had very low levels of HMGB1 with a patchy appearance (**Fig. 4a, b and Extended data Table 1**). Surface associated HMGB1 was lowest in patients with severe inflammation and was absent in areas devoid of IEC (**Fig. 4c**). We assayed FimH in serial tissue sections from the same patients and found higher numbers of FimH positive bacteria in tissue from UC patients (**Fig. 4d, e**) However, FimH was not related to inflammation severity (**Fig. 4f**). When we plotted both surface HMGB1 and FimH from the same patient, UC patients clustered together with low HMGB1 and high FimH, while non-IBD patients clustered together with higher HMGB1 and low FimH (**Fig. 4g**). Mathematically modeling the relationship between HMGB1 and FimH demonstrated that the number of FimH positive bacteria in tissue was dependent on HMGB1 with levels of FimH increasing as HMGB1 decreased (**Fig. 4h**). Thus, UC is characterized by failure of HMGB1 defense with a concomitant increase in tissue-associated bacteria expressing the HMGB1 target protein FimH.

**Figure 4.**
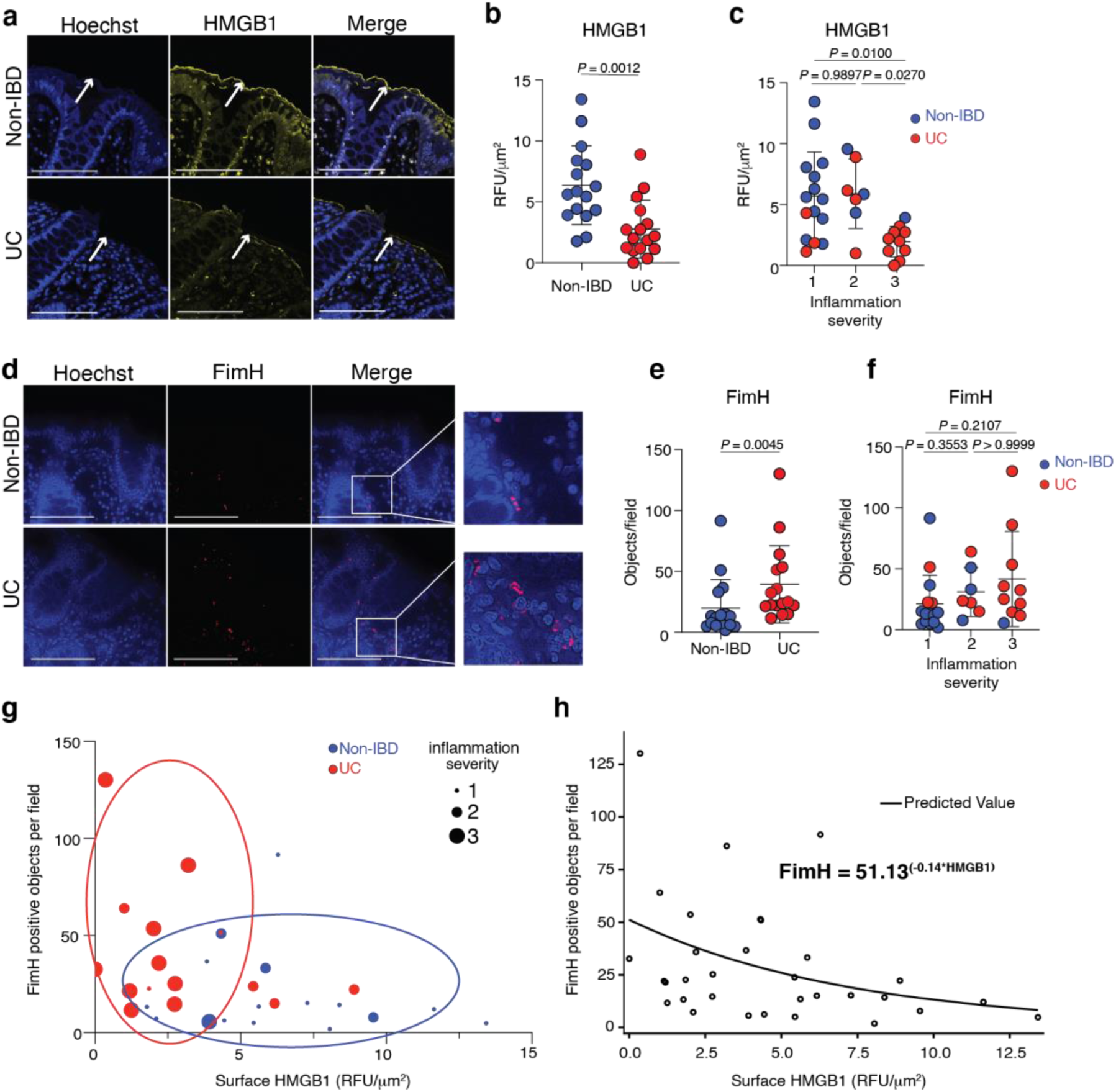
HMGB1 mucosal defense is compromised in ulcerative colitis. **a,** Immunostaining for HMGB1 (yellow) and Hoescht (blue) in Carnoy’s fixed sections of resected colon from Non-IBD or UC patients. (n=16) **b,** Quantification of surface associated HMGB1 using images represented in (a). Staining intensity reported as relative fluorescent units (RFU) per μm^2^. (n=16) **c,** Surface associated HMGB1 reported in (b) graphed by inflammation severity. **d,** Immunostaining for FimH (red) and Hoescht (blue) in Carnoy’s fixed sections of resected colon from Non-IBD or UC patients. Serial tissue sections from the same patients represented in (a) (n=16) **e,** Quantification of FimH positive bacteria using images represented in (d). Reported as number of objects per high powered field. (n=16) **f,** Quantification of FimH positive bacteria reported in (e) graphed by inflammation severity. **g,** Surface HMGB1 and FimH positive bacteria plotted for each patient. The size of the closed circles corresponds to inflammation severity. Open ovals denote the population characteristics by group (non-IBD and UC). **h,** Two-way scatter plot of surface HMGB1 and FimH positive bacteria in each patient with a fitted curve. The relationship between HMGB1 and FimH was captured by non-linear regression. Data are mean ± s.d. Scale bars, 100 µm. Original magnification 400x. Each datapoint represents one individual person. Mann-Whitney U tests were used to compare HMGB1 and FimH in non-IBD vs. UC groups (two-group comparison), and Kruskal Wallis tests were used to assess the difference in HMGB1 and FimH among inflammation groups (three-group comparison).

## Discussion

High mobility group box 1 (HMGB1) is an abundant, ubiquitously expressed protein that has intra- and extracellular functions in many different cell types.^36^ In the gut, HMGB1 is highly expressed in IEC and released into the lamina propria and gut lumen during intestinal infection or when the epithelium is damaged.^37, 38^ HMGB1 in the lamina propria is thought to act as a damage-associated molecular pattern molecule (DAMP) and amplify innate immune responses to microbial signals.^39^ However, no functional role for HMGB1 in the gut lumen has been identified and HMGB1 in the feces is ascribed to passive release from damaged or dying cells.^40–42^ Here we report that HMGB1 is an active component of front-line mucosal barrier defense in the colon with direct and indirect effects to limit virulence of the gut microbiota. Our finding agrees with a previous proteomic survey that detected HMGB1 in mucus isolated by lightly scraping the mouse colon.^43^ It also complements a growing body of literature demonstrating that HMGB1 contributes to both inflammation and antibacterial defense during infection.^44–46^

The cross-kingdom protein-protein interaction between mammalian HMGB1 and bacterial FimH directly limits bacterial virulence by inactivating adhesion through FimH. Expression of T1F genes is also suppressed by HMGB1, suggesting that bacteria like *E. coli* regulate phase or virulence in response to HMGB1. It is possible that the presence of HMGB1 indicates to the bacteria that T1F adhesion will not be effective, or that there is another, as yet unknown effect of HMGB1 on bacterial fitness. Previous reports of HMGB1 effects on *E. coli* viability have been conflicting with one group reporting that HMGB1 kills *E. coli* and another that it does not.^47, 48^ We did not detect killing when *E. coli* were exposed to HMGB1. Instead, in our study, HMGB1 appeared to incapacitate virulence.

The relationship between low levels of HMGB1 and high levels of FimH positive bacteria was noted in both a murine model and in resected colonic tissue from ulcerative colitis patients in our study. Ulcerative colitis has long been linked to adherent microbes, but no single pathogen has been consistently identified across studies.^49, 50^ Our data could provide an explanation for this that is also consistent with the more recent data suggesting that normally commensal bacteria behave like pathogens (i.e. pathobionts) in ulcerative colitis.^51^ We propose that HMGB1, a critical component of host defense, fails in UC patients, allowing bacteria to switch on adhesion mechanisms normally suppressed by HMGB1 and adhere to host tissues. We do not mean to argue that UC is caused by *E. coli*. Rather, we propose that our data provide an explanation for why *E. coli* and potentially other bacteria adhere to intestinal tissue in UC.

The molecular target of HMGB1 in mucosal host defense is a small, evolutionarily conserved amino acid sequence found in adhesins from many different types of bacteria. Much like the ligands that activate pattern recognition receptors, ToH1 appears to be broadly utilized, critical for virulence, and difficult to modify without losing function.^52^ We employed *E. coli* FimH as the exemplar ToH1 positive protein in our mechanistic studies since FimH and T1F adhesion are extremely well characterized and have been implicated in intestinal diseases, including IBD.^27, 29, 53, 54^ The ToH1 sequence is identical and present in FimH from all *E. coli* genomes examined, including *E. coli* that cause infectious diarrheas, strains associated with chronic urinary tract infection, extraintestinal pathogenic *E. coli*, and IBD-associated adherent and invasive *E. coli*. The discovery of ToH1 in FimH has important therapeutic implications since many studies have demonstrated that T1F adhesion is a preferred system for *E. coli* and preventing T1F adhesion has been proposed as a therapeutic strategy for diseases caused by this organism.^23, 27, 55–58^ Bacterial adhesion mechanisms are considered high value targets for nearly all bacterial diseases since blocking adhesion prevents tissue damage, inflammation, and immune activation in disease models and patients.^59, 60^ ToH1 is not restricted to FimH and our experimental data indicate that HMGB1 broadly limits bacterial adhesion, suggesting that our finding may have applications across many different types of bacteria.

Overall, this study demonstrates that HMGB1 actively enforces commensalism in gut bacteria and when it fails, microbes activate virulence and adhere to host tissues. HMGB1 precisely targets bacteria expressing adhesins containing a specific amino acid sequence that is necessary for adhesion to host cells. Therefore, only microbes that have activated virulence are subject to this defense, leaving non-virulent microbes untouched. Our discovery that HMGB1 targets *E. coli* FimH through ToH1 means that failure of this defense is likely relevant for multiple diseases where *E. coli* adheres to the gut epithelium, including infectious diarrheas, colorectal cancer, and IBD. We expect that the same principles of HMGB1 defense will also apply to other bacterial adhesins containing ToH1. However, some bacteria may have additional virulence elements that allow them to evade or degrade HMGB1. Ultimately, our findings represent an important advance toward a functional, rather than taxonomic, understanding of microbial virulence and disease mechanisms. They also provide a novel molecular target for diagnosis and treatment of bacterial disease that has the potential to identify and target only virulence expressing microbes, without regard to their antibiotic susceptibility.

## Extended data figures

**Extended data Figure 1.**
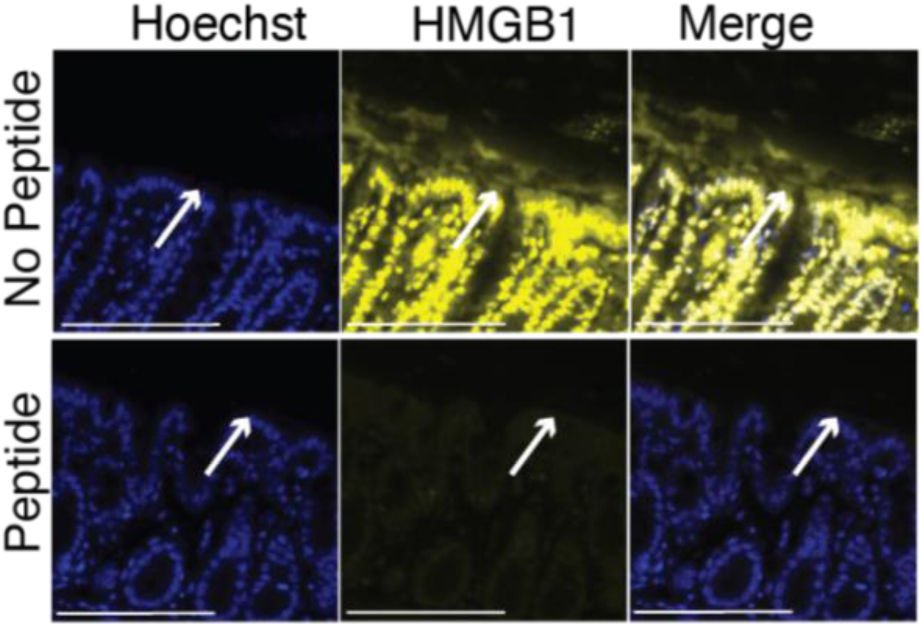
Anti-HMGB1 antibody specifically recognizes HMGB1 in colonic mucus. Immunostaining of Carnoy’s fixed proximal colon sections from HMGB1^WT^ mice either using antibody treated with buffer (No peptide) or antibody incubated with the immunizing peptide (Peptide) to block HMGB1-specific staining. Scale bars, 100 µm. Original magnification 400x.

**Extended data Figure 2.**
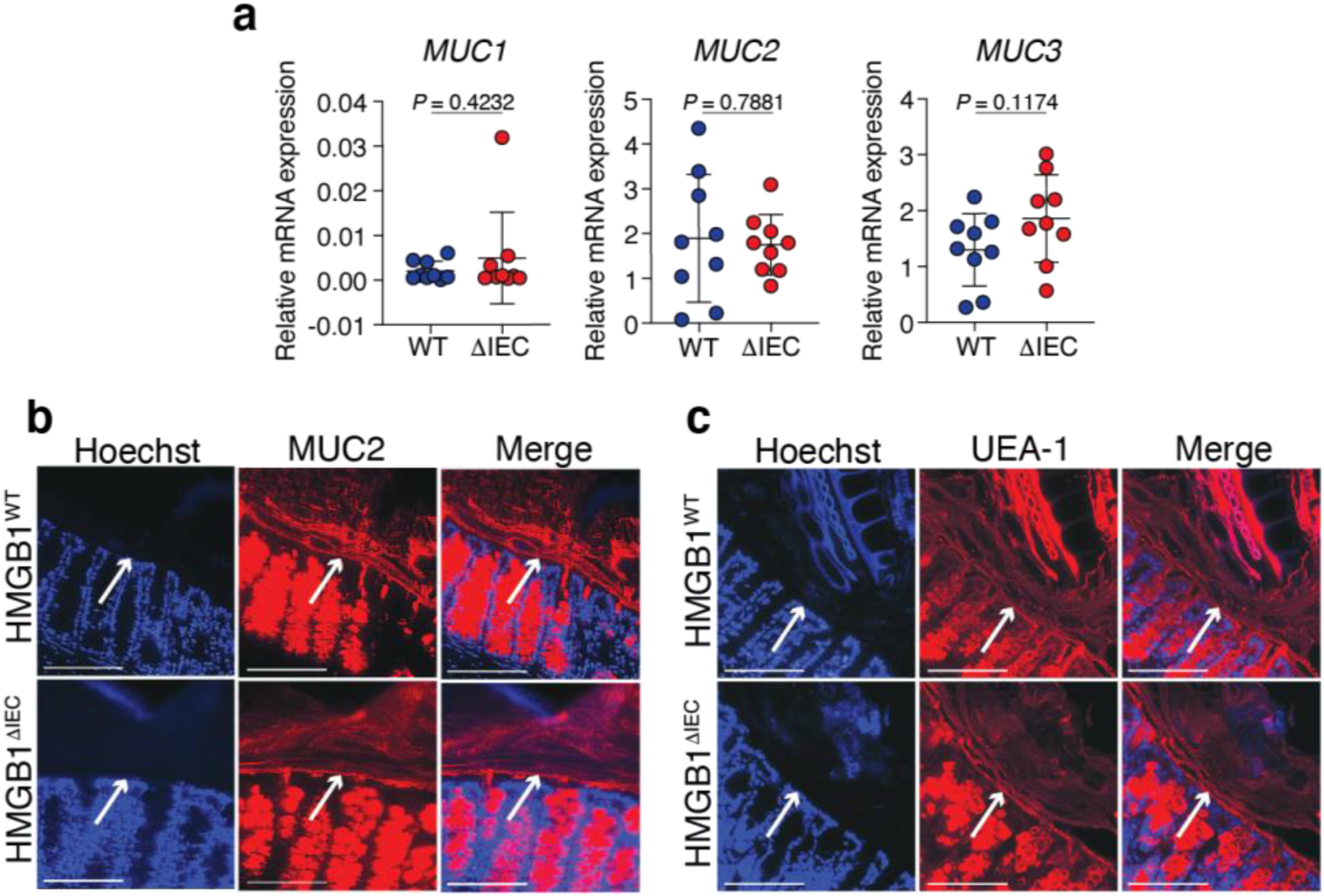
Mucus production is similar in the colons of HMGB1^WT^ and HMGB1^ΔIEC^ mice. **a,** Quantitative reverse transcriptase (qRT) PCR for expression of *Mucin1 (Muc1)*, *Mucin2 (Muc2)*, and *Mucin3 (Muc3)* in colons from HMGB1^WT^ and HMGB1^ΔIEC^ mice. (n=9) **b,** Immunostaining for Mucin 2 (Muc2) in Carnoy’s fixed proximal colon sections from HMGB1^WT^ and HMGB1^ΔIEC^ mice. Arrows indicate the epithelial surface. (n=4) **c,** Lectin staining with fluorescently labeled Ulex europaeus agglutinin-1 (UEA-1) in Carnoy’s fixed proximal colon sections from HMGB1^WT^ and HMGB1^ΔIEC^ mice. Arrows indicate the epithelial surface. (n=4) Data are mean ± s.d. Significance determined by Student’s two-tailed T-tests. Each datapoint represents one individual mouse. Scale bars, 100 µm. Original magnification 400x.

**Extended data Figure 3.**
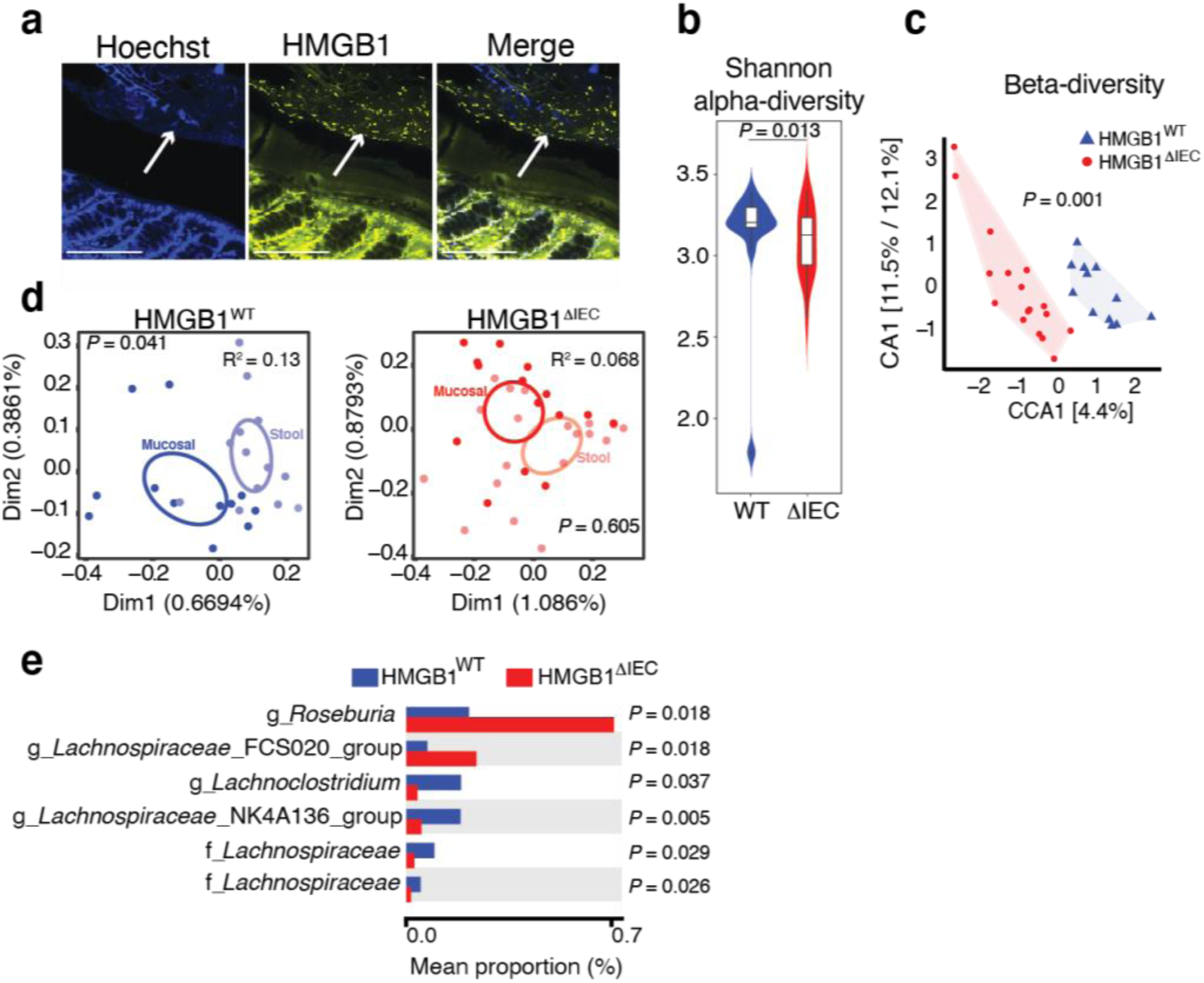
HMGB1 interacts with gut microbes, but does not exert strong selection pressure on the overall microbial community. **a,** Immunostaining for HMGB1 in Carnoy’s fixed proximal colon sections from HMGB1^WT^ mice. Focal plane optimized to capture gut microbes. Scale bars, 100 µm. Original magnification 400x. Arrows indicate the leading edge of gut microbes. (n=20) **b,** Shannon alpha-diversity index of ASV abundances using DNA isolated from mucosal scrapings of colons from HMGB1^WT^ and HMGB1^ΔIEC^ mice. (n=12 WT/18 ΔIEC) **c,** Canonical correspondence analysis (CCA) on ASV abundances using DNA isolated from mucosal scrapings of colons from HMGB1^WT^ and HMGB1^ΔIEC^ mice. (n=12 WT/18 ΔIEC) **d,** Dimensional reduction plots used to characterize microbiome differences between the indicated sites (stool and mucosal) in samples from HMGB1^WT^ and HMGB1^ΔIEC^ mice. (n=12 WT/18 ΔIEC) R^2^ derived from permutational multivariate analysis of variance with site as the main variable. R^2^ indicates the difference between the composition of the microbiota at the two sites in mice of each genotype (HMGB1^WT^ and HMGB1^ΔIEC^) and p-value indicates the significance of the difference in composition between the two sites. (n=12 WT/18 ΔIEC) **e,** Mean proportion of statistically different bacterial strains using DNA isolated from mucosal scrapings of colons from HMGB1^WT^ and HMGB1^ΔIEC^ mice. (n=12 WT/18 ΔIEC) Each datapoint represents one individual mouse except in (d) where each mouse has one datapoint for stool and one for mucosal sample.

**Extended data Figure 4.**
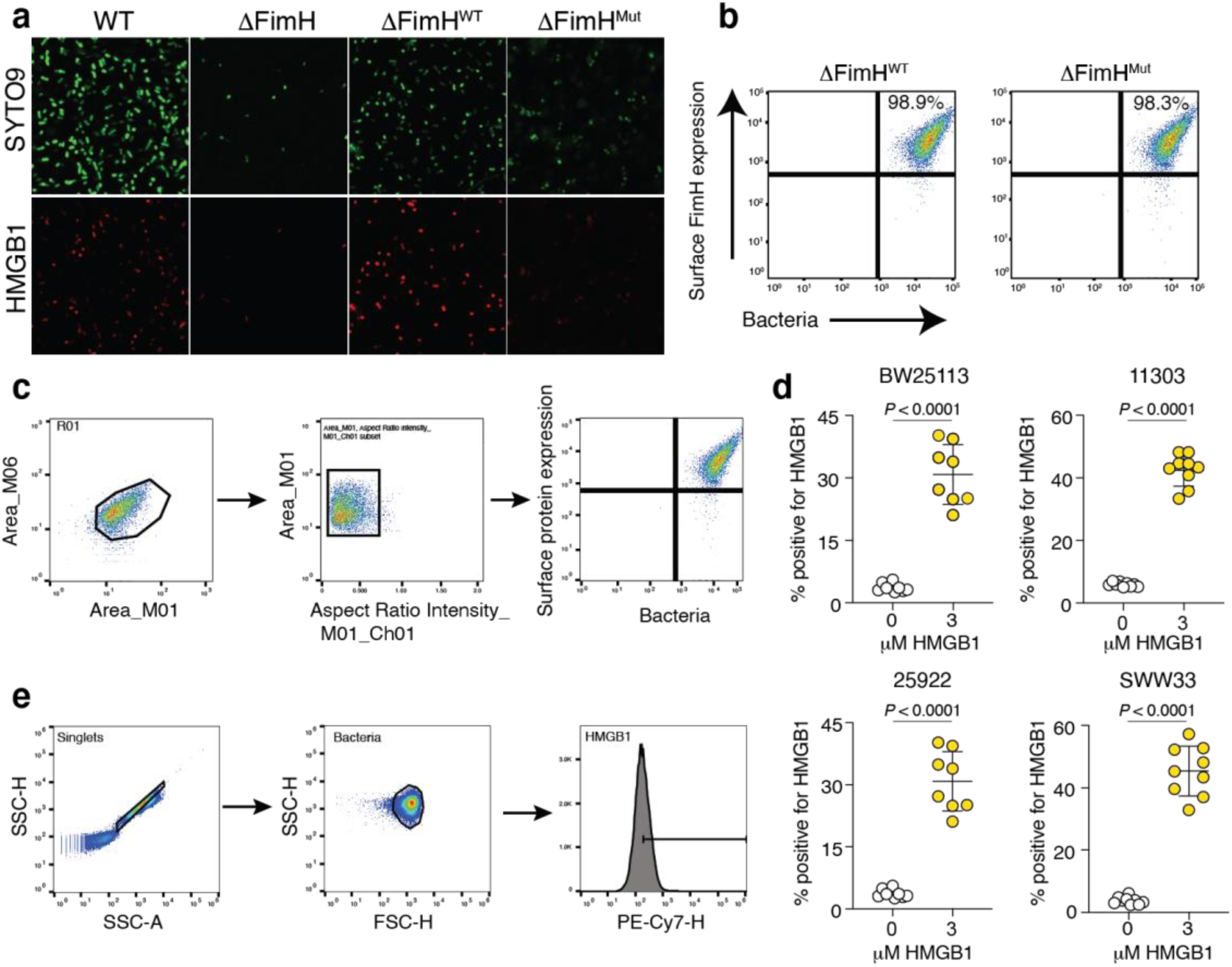
HMGB1 binds to *E. coli* producing FimH containing the ToH1 sequence. **a,** Immunofluorescence staining of HMGB1 (red) bound to *E. coli* (BW25113) (green). Wild type *E. coli* (WT), *E. coli* knocked out for FimH (ΔFimH), or ΔFimH *E. coli* complemented with plasmids carrying either WT FimH (ΔFimH^WT^) or FimH mutated in ToH1 (ΔFimH^Mut^) exposed to rHMGB1 and SYTO 9 to label bacterial DNA. **b,** Flow cytometry for FimH expression on the surface of ΔFimH *E. coli* complemented with plasmids carrying either WT FimH (ΔFimH^WT^) or FimH mutated in ToH1 (ΔFimH^Mut^). **c,** Flow cytometry gating strategy for HMGB1 biding reported in Fig. 3b, c. **d,** Flow cytometry of rHMGB1 binding to *E. coli* of the indicated strains. **e,** Flow cytometry gating strategy for HMGB1 binding reported in (d). Data are mean ± s.d. Significance determined by Student’s two-tailed T-tests. Each datapoint represents one biological replicate.

**Extended data Figure 5.**
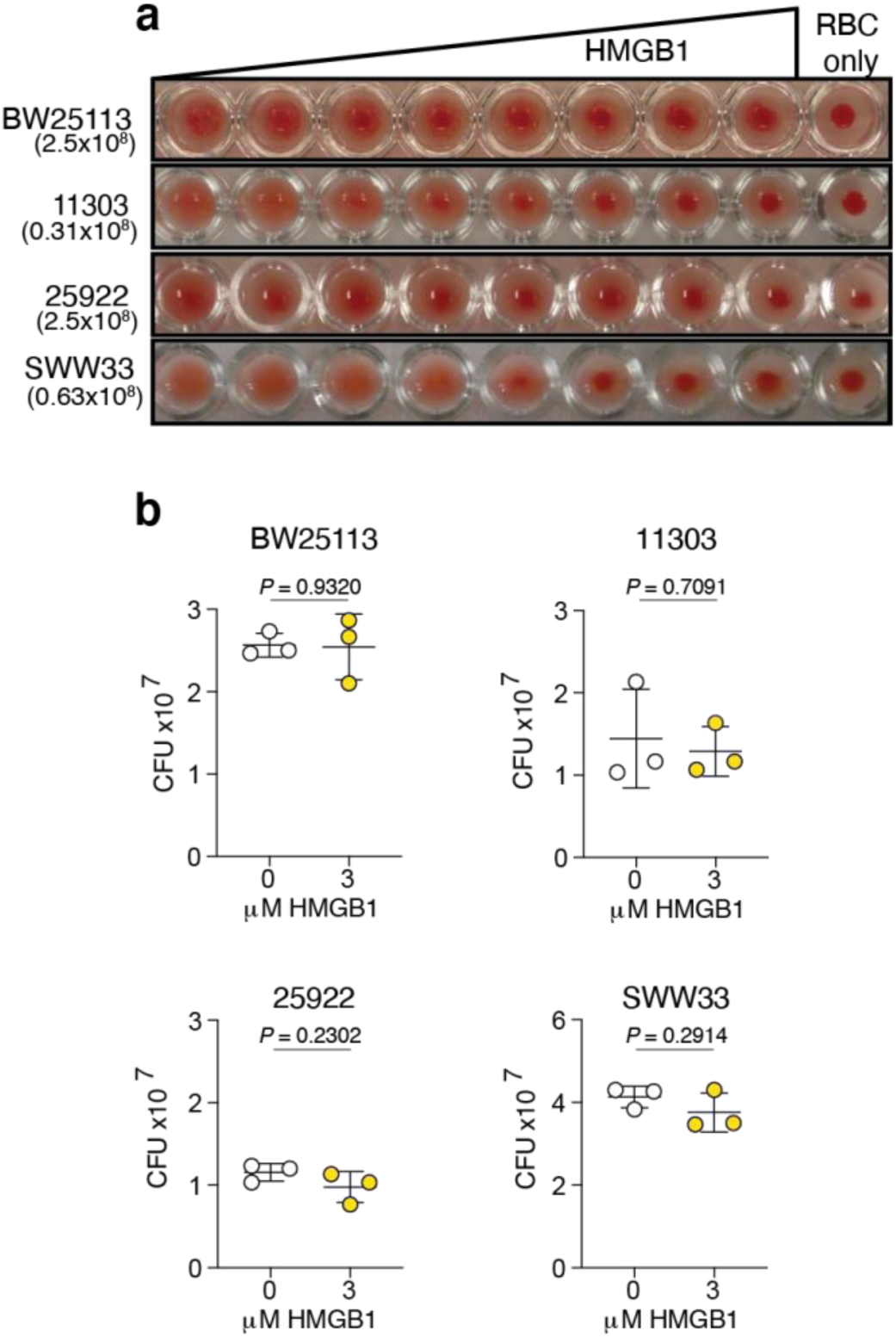
HMGB1 blocks RBC agglutination, but does not kill *E. coli*. **a,** RBC agglutination by various strains of E. coli in the presence of increasing amounts of rHMGB1. Numbers under the strain names indicate the number of bacteria in each well. (3 replicates) **b,** Colony forming units (CFU) of *E. coli* of the indicated strain exposed to buffer (control) or rHMGB1. (n=3; 3 replicates) Data are mean ± s.d. Significance determined by Student’s two-tailed T-tests. Each datapoint represents one biological replicate.

**Extended data Figure 6.**
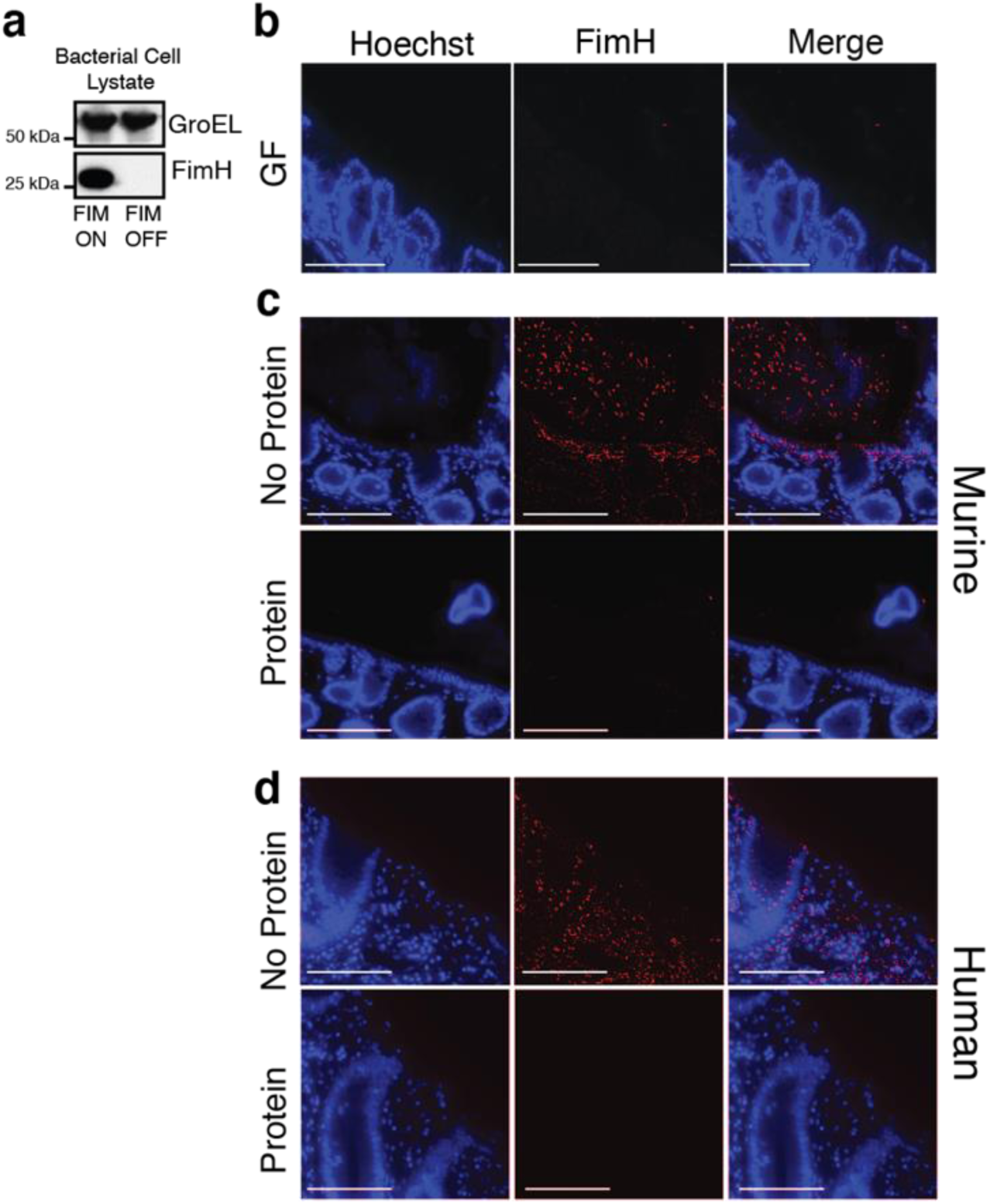
Anti-FimH antibodies specifically recognize FimH. **a,** Immunoblotting using anti-FimH monoclonal antibody on samples of Fim locked *E. coli*. GroEL used as a loading control. **b,** Immunostaining using polyclonal anti-FimH (red) and Hoescht (blue) antibody in Carnoy’s fixed proximal colon sections from GF mice. **c,** Immunostaining using polyclonal anti-FimH (red) antibody and Hoescht (blue) in Carnoy’s fixed proximal colon sections from the mouse and human sample with the highest FimH signal in our studies. Antibody staining without (No Protein) or with (Protein) prior incubation with the immunizing protein to identify signal specific to the immunogen.

**Extended data Table 1.**
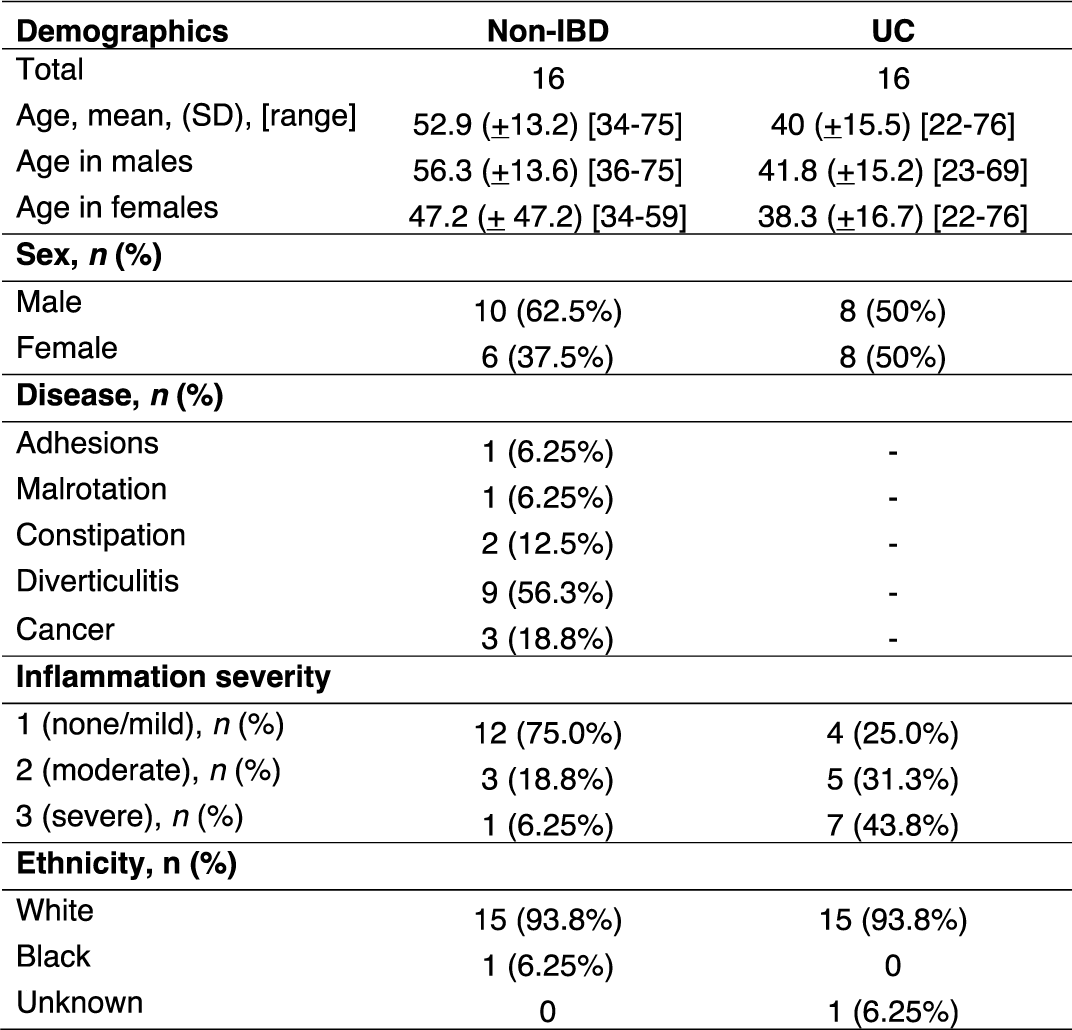
Patient demographic data.

**Extended data Figure 7.**
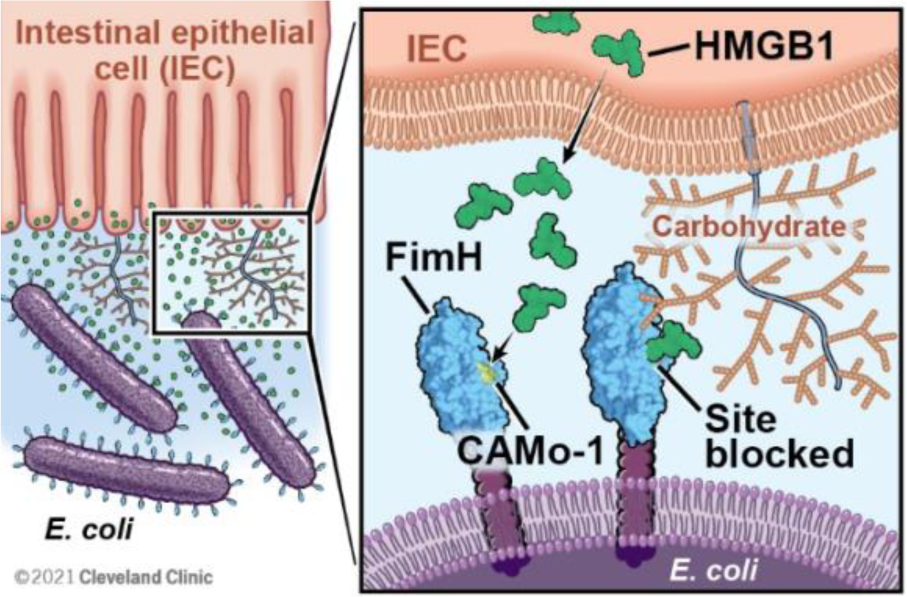
Model of HMGB1 host defense at the gut mucosal barrier. CAMo-1 is the conserved amino acid motif targeted by HMGB1.

## Acknowledgements

We would like to thank Dr. Beat Ernst and Laboratory for the kind gift of plasmids for recombinant production of the FimH lectin domain, Dr. Harry Mobley and Laboratory for the kind gift of T1F phase-locked *E. coli*, Dr. Evgeni Sokurenko and Laboratory for the kind gift of antibodies that recognize FimH, Dr. George Georgiou for the kind gift of HM125 *E. coli*, Dr. Michael Wannemuehler for the kind gift of SWW33 *E. coli*, and Dr. William Schwan for technical assistance. We would also like to thank Dr. Rudi Glockshuber for invaluable technical assistance in producing recombinant FimH. This work utilized resources and equipment provided through the University of Chicago DDRCC (NIDDK P30 DK042086) and the Cleveland DDRCC (NIDDK P30 DK097948). We benefitted from use of the Light Microscopy Core, the Leica SP8 confocal microscope that was purchased with funding from National Institutes of Health SIG grant 1S10OD019972-01, and the Flow Cytometry Core at the Cleveland Clinic Lerner Research Institute. We thank David Schumick for the excellent model illustration of HMGB1 host defense at the gut mucosal barrier. Funding for this study was provided by The American Gastroenterological Association (AGA Microbiome Junior Investigator Research Award to JSM), The Gastrointestinal Research Foundation (JSM), the Falk Medical Research Trust (JSM), and the NIH (1K08DK1114713 and P30 DK42086 to JSM).

## Author contributions

AMCO, BA, MB, XZ, YT, CMC, BM, HS, CF, ZD, and VL planned and performed experiments. NS analyzed microbiome data. DM and AS created the *E. coli* mutants. LX performed statistical analysis. JSM planned and directed the research. AMCO, BA, and JSM wrote the manuscript.

## Materials and correspondence

Jeannette S. Messer

## Methods

### Mice

All animal protocols were approved by the University of Chicago or the Cleveland Clinic Institutional Animal Care and Use Committees. All mice used in this study were on a C57BL/6 genetic background. Specific pathogen-free (SPF) mice were purchased from Jackson Labs. Germ-free (GF) mice were maintained under gnotobiotic conditions at the university of Chicago or Cleveland Clinic prior to euthanasia. Hmgb1*^fl/fl^* (WT) and Hmgb1*^fl/fl,^* ^Vil-CRE^ (ΔIEC) mice were originally created by Eugene Chang’s lab at the University of Chicago as previously described^13^. Mice were maintained under SPF conditions, apart from the GF mice. All animal experiments were performed at least two and in most instances three independent times using both age and sex-matched mice. When possible, littermate controls were used. Mice were between 6 and 12 weeks of age and both sexes were used for all experiments. Mice were housed under a 12-hour light/dark cycle and fed standard laboratory chow.

### Bacteria

#### Culture

All *E. coli* strains were cultured under fimbriae inducing conditions. Strains were passaged twice under static conditions in nutrient broth at 37°C or passaged once on a nutrient agar plate incubated overnight at 37°C unless otherwise indicated. Media consisted of 0.5% Bacto Peptone, 0.3% Beef Extract, 0.5% NaCl and 1.5% agar (if needed), pH 6.8. See extended data table 2 for strain information.

#### BW25113 mutant construction

Full-length FimH was PCR amplified from BW25113 and cloned into the pGex6-1 vector using restriction digest cloning and the BamHI and XhoI restriction sites. Mutation of the ToH1 sequence in FimH was done using two rounds of PCR. Mutation at desired region was introduced in the first round of PCR. Two fragments were generated using primer pairs 1) FimH-For and FimH-Mut1 and 2) FimH-Rev and FimH-Mut2. These two fragments were gel purified, combined, and were used as template for second round of PCR. FimH-For and FimH-Rev were used in the second round of PCR. The products were gel purified, digested by BamHI and XhoI, and cloned into BamHI and XhoI digested pGex6-1 vector. Wild type and mutant plasmids were verified by ability of transformed cells to grow on selection antibiotics and Sanger sequencing. *E. coli* knocked out for FimH (ΔFimH) were transformed with either plasmids containing wild type FimH (ΔFimH^WT^) or FimH mutated in ToH1 ((ΔFimH^Mut^). See extended data table 3 for oligonucleotide information.

#### SWW33 mutant construction

Approximately 1kb homologous recombination arms flanking the sequence to be mutated/deleted (FimH), were PCR amplified from the genomic DNA of WT SWW33. These two arms were then fused together directly or fused to the mutated sequence through PCR. Using TAKARA infusion kit, the PCR-generated fragment was ligated to a suicide plasmid containing kanamycin resistance cassette, R6K origin, RP4 mob, and sacB gene.

This suicide plasmid was transformed into E. coli S17-1λpir through electroporation, and then transformed into *E. coli* SWW33 through conjugation. The successful transconjugants were obtained through plating the conjugation mixture on LB agar plates supplemented with kanamycin and ampicillin. The SWW33 strain that was transformed has an ampicillin resistant plasmid prior to conjugation. The purified transconjugants were then plated on LSW^61^ agar plates supplemented with 10% sucrose to select for the successful knock-in mutants, which were then purified and verified through PCR, Sanger sequencing, and their ability to grow in the presence of ampicillin but not kanamycin. The composition of the LSW agar plates (per liter) was 10 g tryptone, 5 g yeast extract, 5 mL glycerol, 0.4 g NaCl, and 20 g agar. See extended data table 3 for oligonucleotide information.

### Immunostaining

Murine colon tissue was obtained from 8 to12-week-old mice. Human colon samples were residual tissue from colonic resections performed for clinical care of disease (obtained under an existing IRB). Samples with no history of inflammatory bowel disease were used as controls.

All samples were fixed in methyl-Carnoy’s fixative (60% methanol, 30% chloroform,10% glacial acetic acid) for 3 hours at 4°C prior to transfer to 70% ethanol. Paraffin-embedded sections were deparaffinized to water and slides underwent antigen retrieval in 10 mM sodium citrate, 0.05% Tween 20 pH 6.0 using a steamer for 20 minutes and were allowed to cool for 1 hour. They were blocked in serum-free protein block (Agilent, Dako, X0909) and stained with either 10 mg/mL Dylight 594 conjugated Ulex Europaeus Agglutinin I (UEA I), 0.17 mg/mL anti-HMGB1 (Abcam), 0.874 mg/mL anti-FimH polyclonal antibody (custom antibody produced by Genscript) or 0.96 mg/mL anti-Muc2 (Novus). 1mg/mL Alexa Fluor 647 donkey anti-rabbit IgG was used as a secondary if needed. Tyramide SuperBoost (Thermo Fisher Scientific) was used to amplify signals from anti-FimH polyclonal antibody (custom antibody produced by Genscript). Slides were counterstained using 10 mg/mL bisBenzimide H 33258 dissolved in TBS for 20 minutes in the dark at room temperature then cover slipped. For peptide inhibition staining, 0.85 mg/mL of human HMGB1 peptide (Abcam), or 9 ug/mL of full-length FimH protein (antigen for custom antibody production, Genscript) was used.

All immunofluorescent sections were analyzed using either a widefield fluorescent microscope (Keyence BZ-800) or an inverted confocal microscope (40x/1.25 oil objective on a Leica TCS-SP8-AOBS inverted confocal microscope (Leica Microsystems, GmbH, Wetzlar, Germany)).

### Enzyme-linked immunoassay for HMGB1

Samples used for measuring HMGB1 protein levels in the stool and colonic mucus were homogenized in cell lysis buffer (Cell Signaling) containing complete protease inhibitor (Roche) and 100 mM PMSF. After centrifugation at top speed for 15 minutes, supernatants were collected and assayed for protein concentrations via BCA (Thermo Scientific). Samples were analyzed using an HMGB1 detection kit per the manufacturer’s instructions (Chondrex).

### Immunoblot analysis

Bacterial and mucus samples were boiled in Laemmli buffer containing 10% b-mercaptoethanol. All samples were separated using NuPage Bis-Tris gels with MOPS or MES running buffers. Proteins were transferred to PVDF membranes and probed with FimH (Sokurenko Lab), GAPDH (Cell Signaling), 0.17 mg/mL anti-HMGB1 (Abcam), 0.6 mg/mL mouse anti-FimH (Sokurenko (mouse samples) or custom antibody produced by Genscript (bacterial samples), 0.1 mg/mL anti-GroEL (Enzo), or 680-labeled streptavidin overnight in 5% w/v non-fat dry milk Omniblock in 0.1% PBST or LI-COR Intercept PBS blocking buffer at 4°C^62^. After incubation and washing, 0.08 mg/mL of HRP conjugated anti-mouse or anti-rabbit, 0.2 ug/mL 680RD Streptavidin (LI-COR), or 0.2 ug/mL 800RD (LI-COR) anti-mouse secondary antibody was added to the blot and incubated for one hour. After washing, chemiluminescence signal was captured either by film or by using a Bio-Rad ChemiDoc MP and fluorescent signal was detected using an LI-COR Odyssey CLX machine and LI-COR Acquisition software.

### Fluorescent in-situ hybridization (FISH) to detect bacteria

Paraffin-embedded sections were deparaffinized to water. Permeabilization solution (5 mg/mL lysozyme, 0.05 M EDTA, 0.1 M Tris pH 7.4, PBS) was applied to tissue and incubated at 37°C for 20 minutes followed by a PBS wash. Slides were incubated in a hybridization solution (0.9 M NaCl, 0.02 M Tris pH 7.4, 0.005 M EDTA, 1% v/v Triton X-100, 35% deionized formamide, 0.1% w/v BSA, water; pre-warmed to 46°C) for 1 hour in a hybridization oven at 46°C. A universal bacterial probe (EUB338 modified with a 5’ ATTO 647N dye) was denatured in the hybridization solution heated to 95°C for 2 minutes. The slides returned to the hybridization oven covered with 0.1 M probe in hybridization solution for 2.5 hours at 46°C. Slides were washed three times in stringency wash buffer 1 (2x SSC (300 mM NaCl, 30 mM Sodium Citrate, pH 7.0 pre-warmed to 46°C)). A secondary wash was performed three times in stringency wash buffer 2 (0.1x SSC (15 mM NaCl, 1.5 mM Sodium Citrate, pH 7.0)) at room temperature and rinsed in PBS. Tissues were counterstained with 10 mg/mL bisBenzimide H 33258 dissolved in TBS for 20 minutes in the dark at room temperature and coverslipped. The distance between the bacteria and the epithelia was performed using ImageJ analysis software. See extended data table 3 for oligonucleotide information.

### Bacterial flow cytometry

#### HMGB1 binding to E. coli

Bacterial preparations of 3.2 x 10^8^ (O.D. 0.4) *E. coli* were made from an overnight agar culture and incubated with 3 mM HMGB1 recombinant protein containing a HIS tag (Abclonal) or an hFc tag (Sino Biological) and reconstituted in 1mM DTT, 1mM EDTA, PBS buffer for two hours at 37°C. Cells were fixed with 4% PFA and blocked with 1% BSA in PBST for 1 hour. An unconjugated Anti-HIS antibody (Genscript) was added (5.0 mg/mL) and incubated overnight at 4°C. The following day, cells were washed in PBS and secondary antibody (0.25 mg/test PE-Cy7) added for 1 hour at room temperature. Samples incubated with hFc tagged HMGB1 were incubated with PE anti-human IgG Fc recombinant antibody (0.25 mg/test) for 1 hour at room temperature. Finally, cells were stained with 1 mM SYTO9. Bacterial suspensions were analyzed using an Attune NxT flow cytometer (Invitrogen) with FlowJo software (Tree Star).

#### HMGB1 binding to ToH1

Bacterial preparations of 3.2 x 10^8^ (O.D. 0.4) *E. coli* were made from an overnight agar culture and incubated with 3 mM HMGB1(R&D systems) diluted with protein block buffer (Dako) for 2 hours. Bacteria were centrifuged and washed three times with PBS, fixed with 4% PFA for 10 minutes and washed with PBS for three more times. Samples were incubated with anti-FimH (Sokurenko) or rabbit anti-HMGB1 (Abcam) incubated overnight at 4°C^62^. Samples were washed and Invitrogen Alexa a10040, 1:1500 was added. Samples were analyzed using an Image Stream MarkII from Amnis (Luminex).

### Intestinal epithelial cell adhesion assay

Caco2BBe1 cells were seeded onto 0.4 um inserts of a 6.5mm Transwell (Corning) at 250,000 cells per well and allowed to form a polarized monolayer. 72 hours prior to the adhesion assay, 100 ug/mL kifunensine (Cayman Chemicals) was supplemented to 1xDMEM + 10% FBS. Bacterial preparations grown from an overnight nutrient agar culture (1.6 x 10^8^ (O.D. 0.2) *E. coli* per well) were suspended in serum-free 1xDMEM and incubated with 3 mM HMGB1 (R&D), 100 mM mannose (Sigma-Aldrich), or buffer vehicle (1mM DTT, 1mM EDTA, PBS) for 1 hour at room temperature. The apical media from confluent Caco2BBe1 cells were removed and cells washed three times with serum-free 1xDMEM. Treatments were added to wells and cells incubated for 1 hour at 37°C. Cells were washed three times by adding serum-free 1xDMEM and letting the plate shake for 1 minute at 100 RPM between washes. Cells were dislodged with the addition of 0.1% Triton X-100 in PBS, incubated for 5 minutes at room temperature then scraped and collected. Serial dilutions were performed and plated on LB agar plates incubated overnight at 37°C and counted the following day. Serial dilutions of the input *E. coli* were also plated and used to normalize the recovered *E. coli*.

### qRT-PCR

RNA isolation was performed on colonic mucosal scrapings or bacterial lysates using TRIzol (Life Technologies) following the standard extraction procedure. Gene expression was determined using SYBR Green Master Mix (Bio-rad) and a Roche LightCycler 480 System. See extended data table 3 for oligonucleotide information.

### qPCR

DNA isolation was performed on mucosal scrapings or bacterial lysates using DNeasy Powerlyzer Microbial kit per the manufacturer’s instructions. Quantitative PCR was performed using PowerUp SYBR Green Master Mix (Applied Biosystems) in a CFX96 Touch Real-Time PCR system. See extended data table 3 for oligonucleotide information.

### Red blood cell agglutination

Red blood cell agglutination assays were performed as previously described^63^. Briefly, serial dilutions of *E. coli* were incubated with a 3% solution of washed erythrocytes in a round bottom 96 well plate. The wells were mixed by shaking the plate. The plate was incubated at room temperature and imaged 1 hour following shaking. The lowest bacterial number to achieve agglutination was used for the RaMGB1 inhibition assay. For the inhibition assay, *E. coli* were incubated with serial dilutions of HMGB1 recombinant protein (R&D systems) reconstituted in 1mM DTT, 1mM EDTA, PBS buffer and a 3% solution of washed erythrocytes. The wells were mixed by shaking the plate. The plate was incubated at room temperature and imaged 1 hour following shaking.

### Bacterial aggregation Assays

#### E. coli Aggregation

Bacteria expressing green fluorescent protein (GFP) were grown statically under antibiotic selection for two overnight passages before experimental treatment. The final overnight culture was adjusted to an O.D. of 0.4 and 100 mL was centrifuged into a pellet at 5000 RPM for 5 minutes. The supernatants were discarded, and cells were resuspended in TBS with 10 mM CaCl_2_. Bacteria were treated with either 3 mM recombinant HMGB1 (R&D systems) or buffer in a 50 mL reaction at 37°C for 2 hours. Cells were carefully dispensed onto glass coverslips, covered with a 0.15% agarose pad, and imaged using fluorescent microscope. The remaining bacterial samples were analyzed using an LSRFortessa (BD) with FlowJo software (Tree Star) following fixation with 4% PFA.

#### Colonic community microbiota aggregation

Colons from SPF C57BL/6J mice were excised and fileted open. An inoculating loop was used to scrape the contents into a 1.5 mL centrifuge tube containing 1 mL PBS. After centrifugation at 400 G for 5 minutes, the supernatant was strained using a 70 mm cell strainer. The flow through was washed twice with PBS and once with TBS containing 10 mM CaCl_2_ at 10,000 x G for 2 minutes. After the third wash the sample was resuspended in 500 mL TBS/CaCl_2_. The sample was split into 100 uL aliquots and centrifuged at 10,000 x G for 2 minutes. rHMGB1 (R&D systems) was fluorescently labeled with AlexaFluor 647 labeling kit according to the manufacturer’s protocol. Bacteria were resuspended in either 3 mM of labeled rHMGB1 or buffer containing 1 mM SYTO 9 and incubated at 37°C for 2 hours. Cells were carefully dispensed onto glass coverslips, covered with a 0.15% agarose pad, and imaged under fluorescence.

### Recombinant FimH lectin domain production

Recombinant FimH lectin domain protein was produced in HM125 *E. coli* using plasmids provided by the Ernst Laboratory and following described procedures^64^. Purification of the K12 FimH lectin domain was performed as previously described^65^.

### FimH binding to mannose

FimH_LD_ (40 ug/mL) was incubated with serial dilutions of HMGB1 recombinant protein (R&D systems) for 1 hour at room temperature. A mannose coated 96-well plate was washed with 200 uL of 10 mM Tris + 0.05% tween pH 8.0, then the samples were applied to the plate. The plate was sealed and incubated overnight at 4°C. The plate was washed with PBST (0.1%) 4x and blocked with 1% BSA in PBST (0.1%) for 1 hour at room temperature. Then the plate was washed with PBST (0.1%) 4x and the primary anti-HIS antibody was diluted in 0.1% BSA in PBST at a concentration of 5 ug/mL and added to the plate. This was incubated for 1 hour at room temperature. The plate was then washed with PBST (0.1%) 4x and the secondary anti-mouse HRP antibody was added to the plate at 0.16 ug/mL diluted in 0.1% BSA in PBST. The plate was incubated for 30 minutes at room temperature. The plate was washed with PBST (0.1%) 4x and TMB solution was added to the wells. The reaction was quenched with 0.5M sulfuric acid and read at 450 nm on a spectrophotometer.

### *E. coli* killing by HMGB1

Bacterial preparations of 6.4 x 10^8^ (O.D. 0.8) *E. coli* were made from twice passaged nutrient broth cultures and incubated with 3 uM HMGB1 recombinant protein (R&D systems) reconstituted in 1mM DTT, 1mM EDTA, PBS buffer for 2 hours at 37°C. Following incubation, cells were serially diluted, plated onto LB agar plates and colonies were counted the following day after incubation at 37°C.

### FimH expression in *E. coli* exposed to HMGB1

Bacterial preparations of 3.2 x 10^8^ (O.D. 0.4) *E. coli* were made from an overnight agar culture and incubated with 3 μM HMGB1 recombinant protein (R&D systems) reconstituted in 1mM DTT, 1mM EDTA, PBS buffer for 2 hours at 37°C. RNA was isolated using TRIzol (Life Technologies) according to manufacturer’s instructions.

### Fim switch PCR assay

After shaking at 255 rpm for two overnight passages in broth, *E. coli* preparations of 6.4 x 10^8^ cells/mL (O.D. 0.8) were made. The bacteria were treated with 1/100 dilution of conditioned organoid media with (WT) or without (ΔIEC) HMGB1 for 18 hours at 37°C statically. Pilot assays with media derived from small and large intestine derived organoids showed similar results, so additional assays were performed with small intestinal derived organoids since the media contained fewer additives. The conditioned organoid media collected from small intestinal organoids before passaging comprised of Advanced DMEM/F12 supplemented with 1x L-Glutamine (Life Technologies), 10 mM HEPES buffer (Life Technologies), 1x penicillin and streptomycin (Life Technologies), 1x N2 supplement (Life Technologies), 1x B-27 Supplement Minus Vitamin A (Life Technologies), 50 ng/mL murine Epidermal Growth Factor (Peprotech), 100 ng/mL Noggin (Peprotech), 1 μM Jagged 1 (Anaspec), 10 nM Y-27632 (Cayman Chemical Company), and 100 ng/mL R-spondin 1 (Peprotech). This was concentrated using a 3 kDa MWCO filter insert with PBS as the diluent. Bacterial DNA was extracted using DNeasy PowerLyzer Microbial Kit following the manufacturer’s protocol. PCR amplification was performed as previously described^66^. Phase-ON and Phase-OFF fmS PCR product band intensities were standardized against the fsZ amplification product using ImageJ software. See extended data table 3 for oligonucleotide information.

### Quantification of FimH positive bacteria in IF images

Image-Pro Plus 7.0 was used for image operations and measurements. Images were spatially calibrated to units of micrometers and processed as gray values. Background subtraction commensurate with secondary antibody negative controls were applied across the dataset. Semi-automatic counting was performed by manually setting the intensity range overlapped with images. Objects and measurements were collected across five image fields per sample. Measurements of each object were taken to sort and describe post hoc. Two independent people manually applied the intensity ranges with little deviation and similar results. The five image counts were averaged to represent each sample in units of objects per field.

### Quantification of surface associated HMGB1 in IF images

Leica Application Suite (version 3.7.5 or higher) was used for measuring fluorescence intensity in images gathered on a Leica TCS-SP8-AOBS inverted confocal microscope. Regions of interest were drawn around representative surface epithelium and maximized area. Mean fluorescent values (RFU/um^2) were gathered in each image. In background subtraction from the secondary antibody, only negative controls were applied across the dataset. An average of five image values were used to represent each sample. Two independent people identified representative regions of interest (surface epithelium) to collect fluorescent values with similar results.

### Murine mucus isolation

Mucus isolation was performed as previously described^67^.

### Murine mucosal scrape

Colons from mice were excised, fileted open and the contents were washed off. One mL of PBS was added to the colonic tissue in a 1.5. mL centrifuge tube and vortexed vigorously for 2 minutes. The tissue was removed and the mucus containing mixture was centrifuged at top speed for 5 minutes. The semi-transparent mucus layer on top of the dark pellet was removed and resuspended in 1 mL of PBS. 20 mL was saved for BCA. The remainder was centrifuged again and the mucus containing pellet was solubilized in Laemmli buffer for immunoblotting at sample buffer for ELISA.

### Mucus invasion assay

Bacteria expressing GFP were grown statically overnight in broth for two passages and made into preparations of 8 x 10^8^ cells/mL (O.D. 1.0). These *E. coli* preparations were used to fill the middle channel of a chemotaxis µ-Slide (Ibidi). Mucus isolated from WT and DIEC mice were added to opposing reservoirs of the same chamber and incubated at 37°C for 1 hour. Five representative images of fluorescent bacteria invading mucus samples were captured along the leading mucosal edge, proximal to the middle channel of the chamber. These assays were performed at least three times with different bacterial preparations. Any and all image processing settings to reduce background signal were applied across the whole dataset. Measuring the percent relative fluorescence in mucus was performed by using the ImageJ analysis software. After spatially calibrating the image set units to known micrometers, a region of interest was applied beginning at the leading mucosal edge, spanning its full length, and approximately 300 mm deep in each image; approximately totaling 170,000 um^2^; or the maximum area common to all images. Integrated density values were collected and divided by the area of the region of interest (ROI) to account for possible variances upon re-drawing ROIs. Those mean fluorescence values were averaged between the five images and used to represent each sample within the chamber slide.

Given that bacteria had an equal probability of traveling in either well’s direction, the mean fluorescence of each well was compared to the other as a proportion of the total fluorescence of the reaction chamber. Two independent people identified regions of interest to collect fluorescent values with similar results.

### Sulfo-SBED label transfer

The label transfer reaction was performed at previously described^68^. Briefly, rHMGB1 (R&D Biosystems) was treated as the “bait protein” and was reconstituted to 0.2 mg/mL in sterile PBS. Sulfo-SBED was calculated to be added at a 3-molar excess over the purified HMGB1. Sulfo-SBED was dissolved in DMSO to 10 mg/ml and added to the rHMGB1. The reaction was incubated at room temperature for one hour protected from light. Then labeled protein was added to a desalting column (Thermo Fisher #89849) equilibrated with 50 mM HEPES, 150 mM NaCl; pH 7.3 to remove the excess crosslinker and leave the labeled purified HMGB1 in the proper reaction buffer. Sulfo-SBED labeled HMGB1 and unlabeled FimH_LD_ were allowed to incubate together for 2 hours at room temperature protected from light to ensure interaction. The molar ratio used of FimH:HMGB1 was 2.2 mM FimH: 4 mM HMGB1 and was performed in a 50 ml reaction. To crosslink the samples, reactions were moved to a clear 96-well polypropylene low binding plate placed 5 cm from the UV light source (Stratalinker) and exposed to 538 nm wavelengths for 8 minutes on ice. Crosslinked reactions were moved into fresh tubes with the addition of a DTT sample buffer to a final DTT concentration of 100 mM to complete disulfide bond reduction and transfer the biotin label. Samples were heated to 70° C for 10 minutes and analyzed by immunoblot.

### Microbiome studies

#### Microbiome sampling

Littermate mice of both sexes and genotypes were separated into group housing by genotype at weaning. Soiled bedding from mice of both genotypes was collected, mixed, and distributed across all of the cages at the time of separation. At 20 weeks of age, fresh stool samples were collected and the mice were sacrificed for collection of mucosal scrapings. Samples were submitted to the Environmental Sample Preparation and Sequencing Facility at Argonne National Laboratory for analysis of 16S rRNA.

#### Bioinformatics

Individual fastq files without non-biological nucleotides were processed using Divisive Amplicon Denoising Algorithm (DADA) pipeline^69^. The output of the dada2 pipeline (feature table of amplicon sequence variants (an ASV table)) was processed for alpha and beta diversity analysis using phyloseq^70^, and microbiomeSeq (http://www.github.com/umerijaz/microbiomeSeq) packages in R. Alpha diversity estimates were measured within group categories using estimate richness function of the phyloseq package. Multidimensional scaling (MDS, i.e., PCoA) was performed using Bray-Curtis dissimilarity matrix between groups and visualized using the ggplot2 package^71^. As appropriate, we adjusted for multiple comparisons using the BH FDR method while performing multiple testing on taxa abundance across groups^72^. An analysis of variance across the groups for α-diversity was performed. Permutational multivariate analysis of variance (PERMANOVA) was performed on all principal coordinates obtained during PCoA. Linear regression (parametric) and Wilcoxon (non-parametric) were performed on ASV abundances vs metadata variable levels (e.g., diet components) using R base functions.

**Extended data Table 2.** *E. coli* strains used in this study.

| Strain | Source |  |
| --- | --- | --- |
| *Escherichia coli* BW25113 | Coli Genetic Stock Center | https://cgsc.biology.yale.edu/Strain.php?ID=64667 |
| *Escherichia coli* JW4283-3 (ΔFimH) | Coli Genetic Stock Center | https://cgsc.biology.yale.edu/Strain.php?ID=109349 |
| *Escherichia coli* JW4276-1 (ΔFimE) | Coli Genetic Stock Center | https://cgsc.biology.yale.edu/Strain.php?ID=109345 |
| *Escherichia coli* 11303 | ATCC | ATCC 11303 |
| *Escherichia coli* 25922 | ATCC | ATCC 25922 |
| *Escherichia coli* SWW33<br>FimH:TSETPRV | Michael Wannemuehler | Iowa State University |
| *Escherichia coli* SWW33<br>FimH:ASATARA | Mohammed Dwidar | Cleveland Clinic |
| *Escherichia coli* SWW33<br>FimH:AAAAAAA | Mohammed Dwidar | Cleveland Clinic |
| *Escherichia coli* SWW33 (ΔFimH) | Mohammed Dwidar | Cleveland Clinic |
| *Escherichia coli* CFT073 Fim ON | Harry L.T. Mobley | University of Michigan |
| *Escherichia coli* CFT073 Fim OFF | Harry L.T. Mobley | University of Michigan |

**Extended data Table 3.**
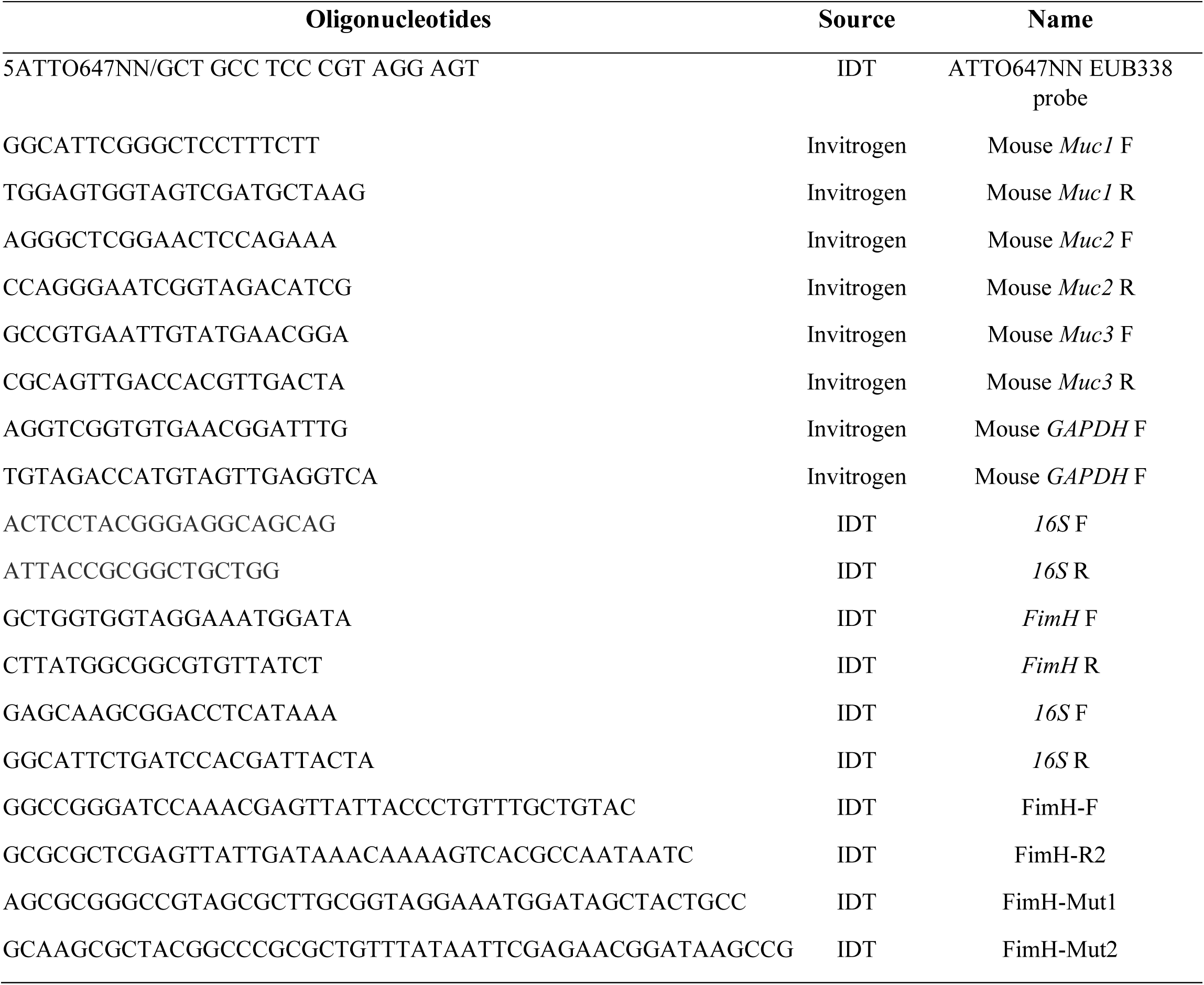
Oligonucleotides used in this study.

